# Analysis of the *FBXO7* promoter reveals overlapping Pax5 and c-Myb binding sites functioning in B cells

**DOI:** 10.1101/792895

**Authors:** Suzanne Randle, Heike Laman

## Abstract

Fbxo7 is a key player in the differentiation and function of numerous blood cell types, and in neurons, oligodendrocytes and spermatocytes. In an effort to gain insight into the physiological and pathological settings where Fbxo7 is likely to play a key role, we sought to define the transcription factors which direct *FBXO7* expression. Using sequence alignments across 28 species, we defined the human *FBXO7* promoter and found that it contains two conserved regions enriched for multiple transcription factor binding sites. Many of these have roles in either neuronal or haematopoietic development. Using various *FBXO7* promoter reporters, we found ELF4, Pax5 and c-Myb have functional binding sites that activate transcription. Overlap of Pax5 and c-Myb binding sites suggest that these factors bind cooperatively to transactivate the *FBXO7* promoter. Although endogenous Pax5 is bound to the *FBXO7* promoter in B cells, c-Myb is also required for *FBXO7* expression. Our data suggest the interplay of multiple transcription factors regulate the *FBXO7* promoter.

## Introduction

F-box proteins (FBP) are exchangeable subunits within <u>S</u>kp1-<u>C</u>ullin1-<u>F</u>-box protein (SCF)-type E3 ubiquitin ligases.^1–3^ These enzymes conjugate a 76aa ubiquitin peptide onto proteins, and this post-translational modification can precipitate that target’s degradation, change of localisation or activity. Fbxo7 is one of ~70 F-box domain-containing proteins, which are receptors for SCF-E3 ligases. However, Fbxo7 also functions outside of canonical ubiquitin-dependent pathways, for example, acting as scaffolds for other regulatory proteins.^4^ At a physiological level, mis-regulation of Fbxo7 has been implicated in human diseases with disparate aetiologies, including Parkinson’s disease, anaemia and cancer, attesting to its critical and pleiotropic role in numerous cell types.

Fbxo7 has hundreds of potential substrates^12^ making the discovery of the critical pathways impacted upon by its mis-regulation in specific pathological contexts challenging. At a molecular level, Fbxo7 affects the cell cycle, increasing cyclin D/Cdk6 activity and stabilising the cyclin-dependent kinase inhibitor, p27;^13^ the regulation of stress-induced mitophagy via the PINK1/Parkin pathway; and NF-κB signalling, via cIAP and TRAF2 interactions,^14^ and BMP signalling via NRAGE-TAK1-TAB1 complex formation.^15;16^ In addition, Fbxo7 ubiquitinates proteasomal subunits, like PSMA2, affecting the assembly of proteasomes,^12;17–19^ and ribosomal subunits, like the stress-responsive ribosomal subunit RPL23, to induce p53 transcriptional responses.^20^ In addition, Fbxo7 is essential for male fertility, and this is attributed to the stabilisation of a proteasomal regulator, PI31.^21–23^ The severity of phenotypes in neurons, erythrocytes, spermatocytes and lymphocytes demonstrate the essentialness of Fbxo7-regulated pathways.

We have shown Fbxo7 has an important role in the development and differentiation of B and T lymphocytes.^24;25^ In addition, Fbxo7 can act as an oncogene as its over-expression in concert with p53 mutation promotes T cell lymphomagenesis.^10^ Recently, a number of high-resolution studies of transcription factor (TF) networks reveal the TFs that function in the normal expansion and differentiation of immature progenitor cells are often dysregulated in leukaemia and lymphoma.^26–34^ Since Fbxo7 and its proto-oncogenic partners, CDK6 and cyclin D2 and D3 are key cell cycle regulators in B and T cells, we investigated whether any of the lineage-specifying haematopoietic TFs control the transcription of Fbxo7. We sought to identify potential TF binding sites within the *FBXO7* promoter and to investigate their regulation of Fbxo7 expression. We identified several promoter elements for TFs, including ELF4, c-Myb and Pax5, which is a master regulator of B cell differentiation, neural development and spermatogenesis.

## Materials and Methods

### Promoter alignment

Transcription start sites (TSS) within the NCBI RefSeq *FBXO7* gene sequence were identified using Eponine software. Fbxo7 orthologues from 28 mammalian species were aligned using the USCS Comparative Genomics 28-way vertebrate alignment and conservation track, and regions of conservation identified. Sequences 10 kb upstream to 10 kb downstream of the gene start site were analysed until conserved regions of similarity stopped. Putative conserved TF binding sites were then identified using MatInspector software. Sites found in more than half of the species were annotated. Data was presented in Clustal W format. This analysis was performed by Dr Michael Mitchell of the CRUK Bioinformatics & Biostatistics Service, who is deceased.

### FBXO7 promoter cloning

A 1.7 kb DNA region containing sequences approximately 1.3 kb upstream to 0.4 kb downstream of the *FBXO7* TSS was amplified by PCR from genomic DNA, and subcloned into pGEM-T Easy vector (Promega). Luciferase reporter constructs were amplified from this plasmid, including a 1.5 kb region (containing the full length *FBXO7* promoter, termed Fbxo7-luc), a 0.5 kb proximal promoter region (proximal Fbxo7-luc), and a 0.6 kb distal promoter region (distal Fbxo7-luc), which were sub-cloned into pTA-luc (Clontech). Site-directed mutagenesis was used to mutate the Pax5 and ETS binding sites. All mutations were verified by sequencing.

### Cell culture and biochemistry

U2OS, Eco Phoenix cells and B cell lines (Nalm6, Ba/F3, Raji, A20) were maintained as in ^25^. Immunoblotting was performed as in ^9^.

### Luciferase assay

U20S cells were transfected with 200 ng of reporter plasmid DNA, 200 ng of pEF-LacZ, and 200 ng of mammalian expression vectors. After 48 hours cells were harvested, and luciferase assays performed as in ^12^.

### RT-qPCR

Experiments were performed as in ^9^. Expression was normalised to cyclophilin levels, and values expressed relative to vector control cells.

### Chromatin immunoprecipitation

Chromatin immunoprecipitation (ChIP) was performed as recommended using the ChIP-IT Express kit (Active Motif). 100 μL of sheared chromatin was immunoprecipitated with 3 μg anti-Pax5, IgG or anti-RNA polymerase II (Human ChIP-IT control kit; Active Motif) and precipitated by magnetic protein G beads.

### Semi-quantitative and quantitative PCR

PCR primer pairs encompassed the putative Pax5 binding site in *FBXO7*, and that previously published for the CD19 promoter,^35^ as well as negative control GAPDH primers supplied in the ChIP-IT control kit. Primer pairs were used in triplicate PCR reactions and performed as in ^9^.

### Expression constructs

Pax5 and ELF4 cDNAs were sub-cloned in frame to FLAG or T7 tags in pcDNA3 (Invitrogen). A Pax5 expression construct, pX-13, was provided by Dr F Baumann-Kubetzko. Full length pCB6-MEF with ELF4 sequences was provided by Dr MA Suico. A Pax5 retroviral construct was generated by sub-cloning into MCSV-IRES-GFP. Pax5 shRNA sequences targeting endogenous mouse Pax5 mRNA were from the RNAi Codex database (codex.cshl.edu). shRNA vectors were cloned into MSCV-LTRmiR30-IRES-GFP. All vectors were verified by sequencing.

### Antibodies and primers

used in this study are listed in Supplementary Information.

## Results

### *The* FBXO7 *promoter contains two TF islands*

To investigate the transcriptional regulation of *FBXO7*, its promoter was analysed for the presence of TF binding sites. Sequence alignment of 28 mammalian species identified a region of conserved sequence similarity upstream of the *FBXO7* transcription start sites (TSS). Consequently, the *FBXO7* promoter was delineated as starting 1300bp upstream from the start of exon 1, to 100bp downstream, numbered according to the human *FBXO7* gene (−1300 to +100). Within this region, two clusters of conserved TF binding sites were identified, approximately 1kb apart. One island in the distal promoter region was 125bp in length (−1275 to −1150), while the other in the proximal promoter region was 400bp in length and overlapped the TSS and start of exon 1 (−300 to +100). These islands contained the majority of TF binding sites, and selected regions from 13 of the species surveyed are shown in **Figure 1A**. 32 putative binding sites were identified for 24 different TFs (17 in the distal region; 15 in the proximal region). We identified ‘core’ promoter elements, including two CCAAT boxes on opposite strands at −1257 and −1198 in the distal region, and a GC box, identified as a Sp1 binding site, in the proximal region within exon 1 (+81). No TATA box was identified, and the proximal region was GC rich, suggesting the *FBXO7* promoter is associated with a CpG island. In addition to ubiquitously expressed TFs, like activator protein (AP) and E2Fs, we identified multiple consensus sites for TFs with roles in haematopoiesis and neuronal development (ETS, c-Myb, DMTF, Pax5, Myt1, NeuroD, NRF1, CLOX, ZF5F).

**Figure 1:**
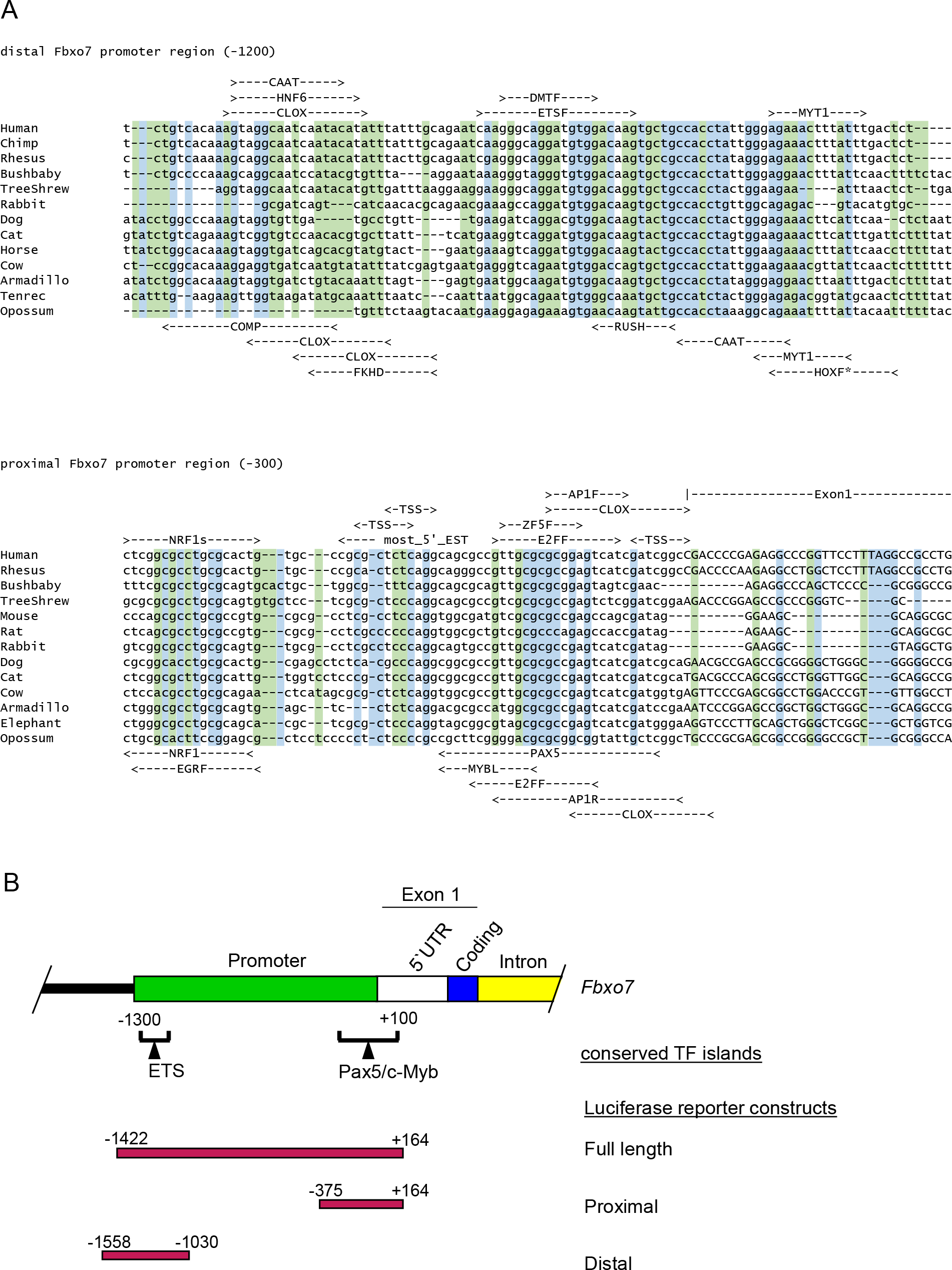
Sequence alignment of 13 mammalian species (**A**, upper) showing −1275 to −1176 of the distal TF island which covers −1275 to −1151, and (**A**, lower) from −66 to +35 of the proximal TF island which covers −300 to +100, numbered according to the human sequence from the start of exon 1. Putative TF binding sites are listed above the alignment on the forward strand, and below the alignment on the reverse strand. Conserved bases are shown in blue and conserved pyrimidines or purines, in green. (**B**) Schematic of the start of human *FBXO7*. The promoter (green) was defined by sequence alignment of several species (see M&M). Several TF binding sites were identified within conserved TF islands (brackets). The 5’ UTR (white), coding sequences (blue), and first intron (yellow) are indicated. Luciferase reporters (red) are below.

### *The* FBXO7 *reporter is activated by ETS factors, including ELF4*

To test putative binding sites, *FBXO7* reporters were generated. Full length *FBXO7* promoter (−1422 to +164), as well as either the distal (−1558 to −1030) or proximal (−375 to +164) regions were cloned into pTA-luc luciferase plasmid (**Figure 1B**). Within the distal region, we identified two ETS binding sites (EBS), shown in **Figure 1A** as ETSF. The consensus site contains a core GGAA sequence, but flanking sequences and co-factor binding impart specificity for particular ETS factors. We tested a panel of ETS factors (ETS1, ETS2, Fli1, ELF4, ELF1, and PU.1) by co-transfecting them with the distal reporter (Distal Fbxo7-luc) or a control (pTA-luc) into U20S cells (**Figure 2A**). We found ETS1, ETS2, ELF1 and ELF4 showed significant activation of the reporter (p<0.05), indicating the EBS were functional. We noted both EBSs contain a WGGA (where W is A/T) sequence, which matches the consensus site for ELF4, consistent with its higher activation of the reporter.^36^ Mutation of the EBSs significantly reduced ELF4 transactivation (p<0.05) (**Figure 2B**), indicating that ELF4 utilised these sites to activate the distal reporter.

**Figure 2:**
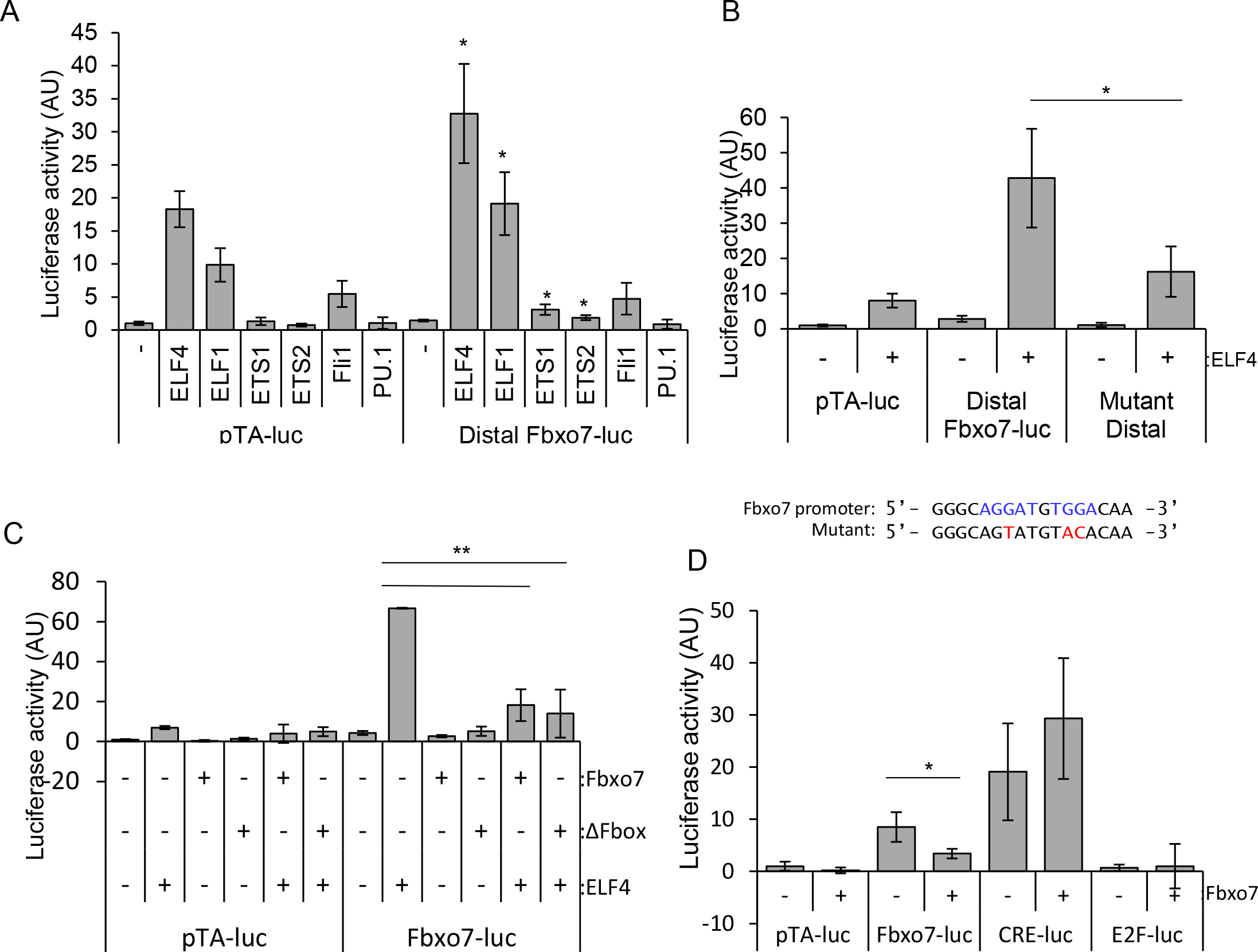
(**A**) Luciferase assay of cell lysates from U2OS cells transfected with distal Fbxo7-luc, or pTA-luc empty vector, along with a panel of ETS family members. Luciferase values in triplicate, were background corrected with non-transfected cell lysate values, and normalised to co-transfected b-galactosidase levels, and expressed relative to empty vector. \**p*<0.05 compared to the relevant empty control levels, n=3. (**B**) Luciferase assay showing ELF4 activation of the empty vector, distal Fbxo7-luc and mutated distal Fbxo7-luc reporter constructs, *n*=3. Below: the two ELF4 consensus sites in the *FBXO7* promoter are shown in blue, and mutated base pairs in red. (**C**) Luciferase assays showing ELF4 activity on the Fbxo7-luc reporter in the presence of exogenous Fbxo7 or ubiquitination dead mutant Fbxo7-ΔFbox. (**D**) Luciferase assays in cells transfected with a panel of luciferase reporters with or without Fbxo7, *n*=3.

### Fbxo7 inhibits ELF4 trans-activation

Lui and co-workers reported Fbxo7 and ELF4 physically interact,^37^ which suggests Fbxo7 may affect ELF4 transactivation. To test this, ELF4 was co-transfected with either WT Fbxo7 or a mutant lacking the F-box domain (ΔF-box), and the full-length reporter into cells. ELF4 activation of the reporter was inhibited by 80% by the addition of WT or mutant Fbxo7 (**Figure 2C**), indicating Fbxo7 inhibits ELF4 transactivation in a ubiquitin-independent manner. We next tested whether this effect was dependent on ELF4 or a general effect on *FBXO7* and other reporters, by transfecting Fbxo7 and measuring luciferase expression from other reporters. As with the distal reporter, we found Fbxo7 significantly inhibited by 50% the full length Fbxo7-, but not the Cre- or E2F-luciferase reporters (**Figure 4D**). Moreover, as U20S cells do not express ELF4, it suggests that Fbxo7 may bind to other ETS factors. Together, these data suggest Fbxo7 can inhibit transcription from its own promoter in an SCF-independent manner.

### *An* FBXO7 *gene reporter is activated by Pax5 and c-Myb*

ETS factors act in concert with other TFs, like Pax5. We identified a Pax5 consensus site in the proximal region. To test its functionality, U20S cells were co-transfected with a Pax5 expression construct or a control and the reporters, and luciferase assays were performed. A dose-dependent increase in luciferase activity was detected with increasing Pax5 expression from the full length and proximal *FBXO7* constructs (**Figure 3A**), with a more limited response from the distal reporter. This suggests that Pax5 activation of the full-length promoter is predominantly via the proximal region. We also mutated the Pax5 binding site within the proximal reporter (**Figure 3B**) and tested the effect (**Figure 3C**). Expression of Pax5 resulted in a 40-fold induction of luciferase activity from the proximal reporter, whereas Pax5 activation of the mutant reporter was significantly reduced by 37% (**Figure 3C**). These data argue Pax5 activation of the reporter occurs at the Pax5 binding site. We next tested whether Pax5 activation was modulated by ETS family members. As in **Fig 3A**, Pax5 activated the full-length reporter (**Figure 3D**). However, when Fli1 was co-transfected with Pax5, luciferase levels increased by 70% over Pax5 alone, and 340% more than Fli1 alone. In contrast, addition of ETS1 or PU.1 inhibited Pax5 activation by 40% and 60%, respectively.

**Figure 3:**
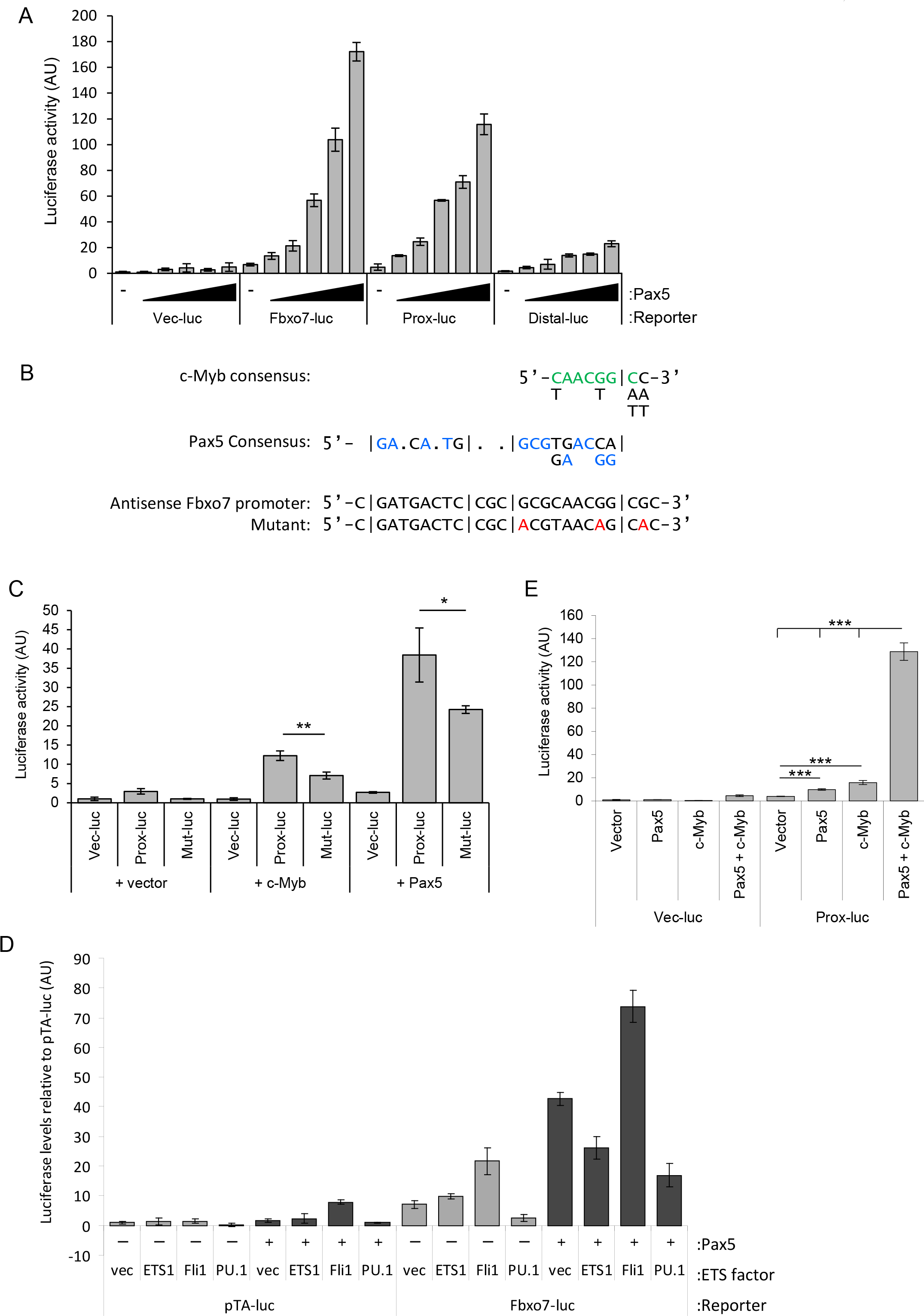
(**A**) Luciferase assay with various *FBXO7* reporters and increasing doses of Pax5. (**B**) A c-Myb binding site overlaps a Pax5 binding site. Shown are the Pax5 binding site (blue), with bases in the Fbxo7 promoter matching the consensus for c-Myb (green), and the mutated bases in the Proximal *FBXO7* reporter (red). (**C**) Luciferase assay on U2OS cell lysates co-transfected with Pax5 or c-Myb, and the empty reporter construct, the WT or mutated proximal Fbxo7-luc. (**D**) Luciferase assay showing activation of various Fbxo7 luciferase vectors by vector only (vec) or exogenous ETS family members, with (+, dark grey) or without (−, light gray) Pax5. (**E**) Luciferase assay on U2OS cell lysates co-transfected with Pax5 or c-Myb, in combination, with the proximal Fbxo7-luc reporter.

**Figure 4:**
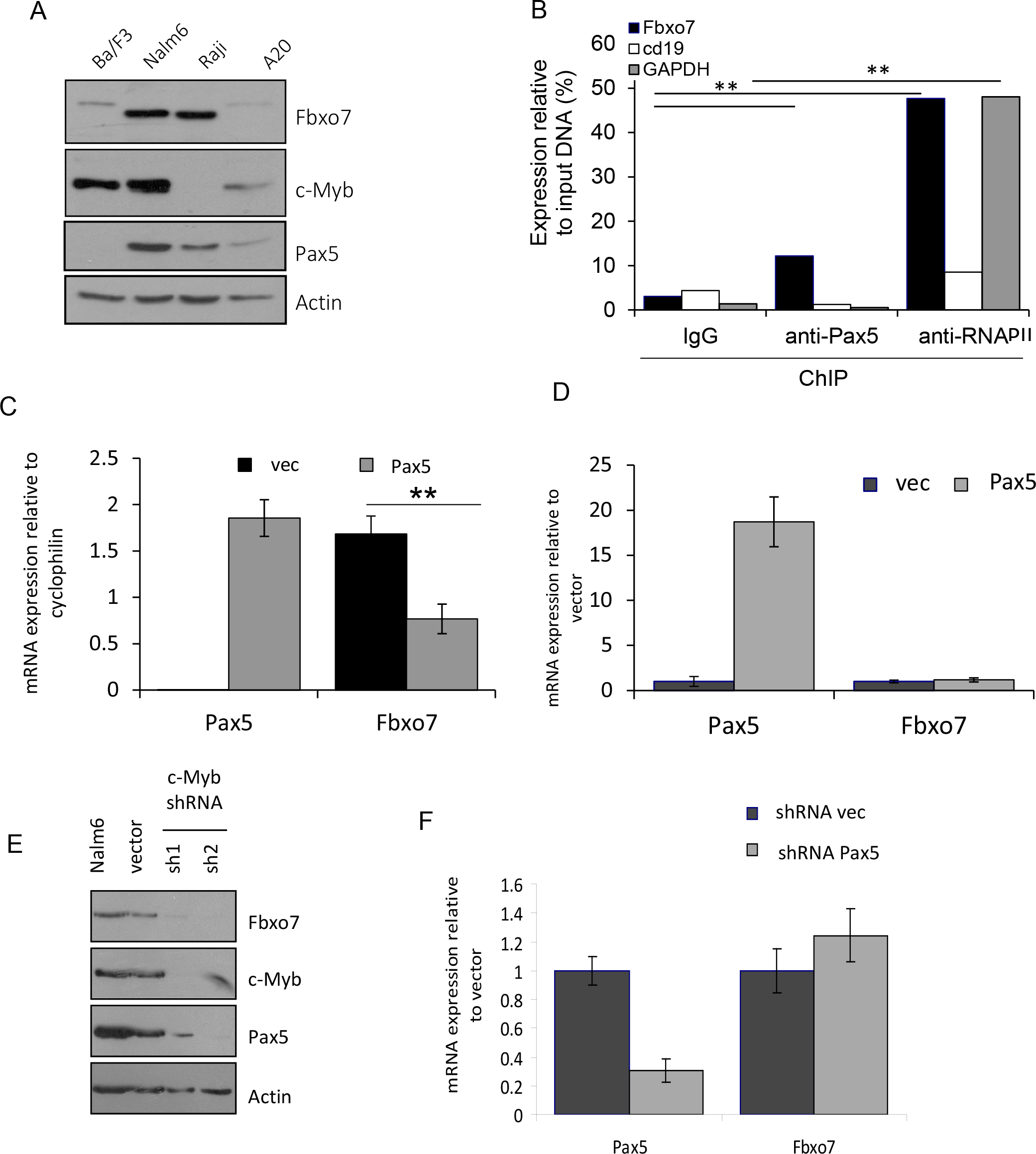
(**A**) B cell lines at various differentiation stages analysed by immunoblotting for the expression of Fbxo7 and Pax5. (**B**) Pax5 chromatin immunoprecipitation of the *FBXO7* promoter in Nalm6 cells. Values were then expressed as a percentage relative to input. \*\**p*<0.01 compared to relevant control IgG levels. *n*=2 independent experiments with triplicate qPCR reactions. Graphs of mRNA expression of Ba/F3 cells (**C**) or Nalm6 cells (**D**) transduced to over-express Pax5 or empty vector, *n*=2. (**E**) Immunoblotting of Nalm6 cell lysates transduced with shRNA constructs targeting c-Myb expression, *n*=2. (**F**) Graph of mRNA expression in Nalm6 cells transduced to express a shRNA construct targeting Pax5, or empty vector. Pax5 and Fbxo7 mRNA levels were analysed by qRT-PCR, normalised to cyclophilin A levels and expressed relative to empty vector control cells.

A c-Myb binding site overlaps the Pax5 binding site (**Figure 1A**), and c-Myb has also been shown to influence Pax5 activity.^38–40^ To test the c-Myb binding site, we transfected c-Myb and the WT and mutated proximal reporters (**Figure 3B**) and found c-Myb activated the proximal region by 12-fold (**Figure 3C**). Furthermore, the mutated reporter, which contained one altered base pair within the consensus c-Myb site, showed decreased activation with c-Myb by 40% compared to the WT promoter. As Pax5 and c-Myb both activated the proximal reporter through an overlapping binding site, we tested whether they had a synergistic or competitive effect on transactivation. When expressed together, Pax5 and c-Myb activated the proximal reporter more than 8-fold over c-Myb alone, and over 13-fold than Pax5 alone (**Figure 3E**) suggesting they act synergistically through this site. These data indicate that Pax5 together with ETS and c-Myb TFs transactivate the *FBXO7* reporters.

### Endogenous Pax5 bound the Fbxo7 promoter

As Pax5 and c-Myb are both involved in B cell development, a screen of c-Myb, Pax5 and Fbxo7 protein expression in B cell lines at different stages of maturation was conducted to determine any correlation (**Figure 4A**). Immunoblotting of B cell lines from progenitor to mature B cell stages showed c-Myb expression was highest in earlier stage pro-B cells (Ba/F3) and pre-B cells (Nalm6), in line with its key role in pro-B cell differentiation. Pax5 expression was highest in pre-B cells, and declined as B cells matured, as expected since downregulation of Pax5 promotes plasma cell differentiation. Fbxo7 expression correlated with greater levels of Pax5, but not necessarily c-Myb, with highest Fbxo7 expression evident in pre-B cells (Nalm6) and B cells (Raji), consistent with Fbxo7’s role in promoting pro-B cell differentiation.^25^

We tested whether Pax5 was present at the *Fbxo7* promoter in maturing B cells, using chromatin immunoprecipitation (ChIP) assays. The *CD19* promoter, a Pax5 target, and *GAPDH*, which does not contain a consensus Pax5 site, were selected as promoter controls, and antibodies to RNA polymerase II and normal serum IgG were used as immunoprecipitation controls. ChIP experiments were performed in Nalm6 pre-B cells, which showed the highest expression of Pax5, Fbxo7, and c-Myb. Immunoprecipitated DNA was analysed by qPCR, and DNA enrichment expressed as a percentage of input DNA to normalize for differences in PCR efficiency. The *FBXO7* promoter was enriched 4-fold in DNA immunoprecipitated using Pax5 antibody compared to IgG only, and 32-fold over the amount of *GAPDH* promoter region immunoprecipitated by Pax5 antibody (**Figure 4B**). This was despite a significant enrichment of the *GAPDH* promoter by RNA polymerase II antibodies. Enrichment of the *FBXO7* promoter in Pax5 immunoprecipitates, and enrichment of the *FBXO7* and *GAPDH* promoter in RNA polymerase II immunoprecipitates were significantly increased (p<0.01) compared to IgG levels. Unexpectedly, there was no significant enrichment of the *CD19* promoter in Pax5 or RNA polymerase II immunoprecipitates, suggesting CD19 was not transcribed. We confirmed the lack of CD19 expression by flow cytometry of the cells with a fluorescently labelled anti-CD19 antibody (data not shown). Similar enrichment of the *FBXO7* promoter in Pax5 immunoprecipitates were found in murine A20 cells. These data indicate the Pax5 binding site in the *FBXO7* promoter is a functional TF binding site in B cells and suggest Pax5 directly regulates *FBXO7* expression.

### Early expression of Pax5 reduced Fbxo7 expression in pro-B cells but not in later stage pre-B cells

We next tested whether earlier expression of Pax5 in Ba/F3 (pro-B) would affect Fbxo7 expression. Ba/F3 cells were infected with retroviruses expressing Pax5 and GFP, RNA was harvested from GFP+ cells, and mRNA levels were assayed using RT-qPCR. We observed a 50% decrease in Fbxo7 mRNA levels when Pax5 expression was introduced (**Figure 4C**). However, no changes in Fbxo7 mRNA levels were seen upon increasing Pax5 expression in Nalm6 or A20 cells which both express endogenous Pax5 (**Figure 4D**). Thus early expression of Pax5 represses *FBXO7* transcription.

As c-Myb is expressed in earlier stage B cells, we tested the effect of reducing its expression. Nalm6 cells were infected with two shRNA constructs targeting c-Myb, GFP+ cells were lysed and assayed by immunoblotting (**Figure 4E**). We observed a strong reduction in Fbxo7 in cells with reduced c-Myb expression, and surprisingly, a concomitant reduction in Pax5 expression. To distinguish whether reduced Pax5 or c-Myb levels decreased Fbxo7 expression, we reduced Pax5 expression using a short hairpin construct in Nalm6 cells; however, there was no discernible change in Fbxo7 expression (**Figure 4F**). Our data are consistent with Pax5 and c-Myb cooperating in activating endogenous Fbxo7 expression at the pre-B cell stage of differentiation.

## Discussion

To investigate Fbxo7 transcriptional regulation, we defined the human *FBXO7* promoter, as a conserved promoter region between −1300 and +100 bp from the start of exon 1. Within the promoter two conserved TF islands (−300 to +100 and −1275 to −1150) were identified. A similar proximal promoter was reported in the pig, although it was limited to 1000 bp upstream of the TSS.^41^ Although no TATA box was identified, as for the pig, the CCAAT and GC boxes may constitute part of the core promoter of *FBXO7*. Consistent with this, analysis of the publicly available gene annotation database Encyclopaedia of DNA Elements (ENCODE; http://genome.ucsc.edu/ENCODE/)^42^, suggests that the proximal region, along with exon 1 of *FBXO7* lies within a CpG island, H3K4 tri-methylation region and DNase I hypersensitivity site, indicating these sequence form the core *FBXO7* promoter.

Given the key roles of the Fbxo7 in blood, we investigated TF binding sites with known roles in haematopoiesis. These included twin ETS binding sites in the distal region of *FBXO7* and overlapping Pax5 and c-Myb sites in the proximal region. We found Pax5, c-Myb and the ETS family members, ELF4 and ELF1, were the strongest activators of a synthetic *FBXO7* reporter. Interestingly, Fbxo7 has been previously reported to directly interact with ELF4. Our data indicates that Fbxo7 inhibits ELF4 activation of the *FBXO7* reporter, an activity independent of its ubiquitin ligase function, suggesting a negative feedback mechanism. Also, since Fbxo7 did not inhibit other luciferase reporters, this suggests Fbxo7 specifically inhibits its own transcription.

The *FBXO7* reporter was also transactivated by Pax5, whose expression is largely restricted to B cells. Using ChIP assays, we found that endogenous Pax5 was bound to the proximal region of *FBXO7* in both the mature B cell line, A20, and a leukaemic pre-B cell line, Nalm6, indicating a functional Pax5 binding site. Despite this, only the aberrant expression of Pax5 in Ba/F3 cells caused a change in *FBXO7* mRNA expression, repressing transcription. It is known that Pax5 acts in concert with other TFs such as ETS factors.^38;43;44^ Pax5 recruits ETS factors, Net and Elk-1, to the mb-1 promoter in pre-B cells to increase DNA binding,^44^ whereas PU.1 recruits Grg4 to Pax5-occupied promoters where Pax5 represses transcription.^43^ We found several ETS factors modulated Pax5 transactivation of an *FBXO7* reporter, including Fli1 which increased transcription, and ETS1 and PU.1 which inhibited it. Interestingly, a study of TF networks in haematopoietic cells using combined data from 53 ChIP-seq studies, identified PU.1 at the *FBXO7* promoter, highlighting that PU.1 also regulates *FBXO7* expression.^45^ Pax5 also enlists other TFs like c-Myb, e.g., Pax5 recruits c-Myb to the RAG-2 promoter in B cells,^38^ and our data indicate Pax5 and c-Myb work cooperatively at the transcriptional start site of the *FBXO7* promoter.

Although Pax5, c-Myb, ETS, have important roles in haematopoiesis, c-Myb and Pax5 also have roles in neural progenitor cell proliferation^46^ and early midbrain development,^47^ respectively. Fbxo7 is expressed in mouse adult brain,^20^ so these potential TF sites active in brain development warrant further study. Other putative regulatory TFs for Fbxo7 had roles in brain development, like neurogenic differentiation factor 1 (NeuroD), which regulates neuronal differentiation fate,^48^ and myelin transcription factor 1 (Myt1), which modulates oligodendrocyte proliferation and differentiation.^49^ Intriguingly, given its role in Parkinson’s disease, a site for nuclear respiratory factor 1 (NRF1) was also found. This TF is associated with mitochondria function and metabolism, neurite outgrowth, and the ‘bounce back’ of proteasome transcription under proteotoxic stress.^50–57^ That multiple TFs co-operate regulate Fbxo7 expression, and are involved in cell differentiation and also stress-responsive functions points to a model wherein the proteins that create and specify particular cellular lineages and mature cell types also have a role in maintaining their cell health when they come under stress.

## Acknowledgements

This work was supported by the BBSRC (BB/J007846/1) to HL.

## Supplementary Material

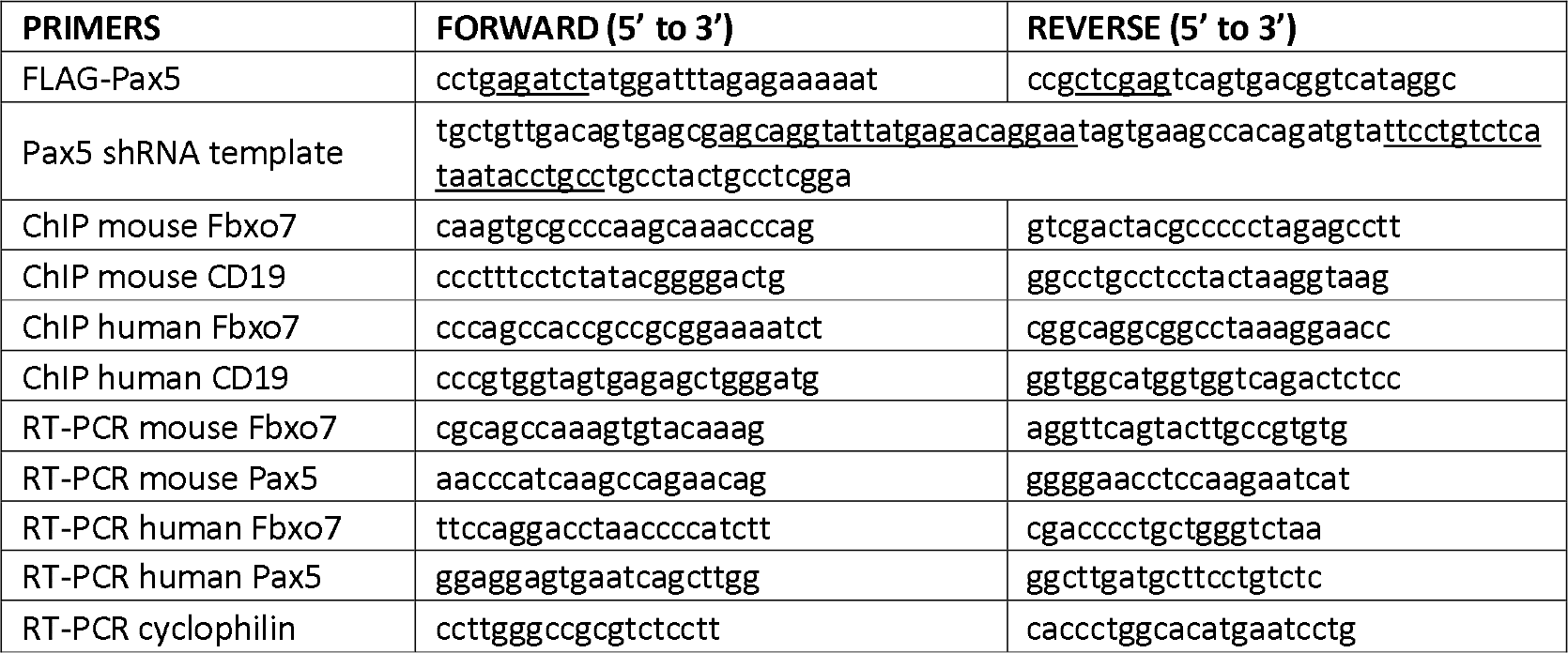

| ANTIBODY | SPECIES | DILUTION | SOURCE | CATALOG # |
| --- | --- | --- | --- | --- |
| Actin | Rabbit | 1:5000 | Sigma | A2066 |
| Fbxo7 (Ab2) * | Rabbit | 1:500 | In house | C-terminus |
| Fbxo7 (Ab1) | Rabbit | 1:1000 | Aviva | ARP43128_P050 |
| Pax5 | Mouse | 1:500 | Santa Cruz | sc-55515 |
| c-myb | Mouse | 1:1000 | Millipore | 05-175 |
| Anti-Mouse | Goat | 1:10000 | Santa Cruz | sc-2055 |
| Anti-Rabbit | Donkey | 1:10000 | Santa Cruz | sc-2313 |

## References

1. J. T. Winston, D. M. Koepp, C. Zhu, S. J. Elledge, J. W. Harper, Curr.Biol. 9, 1180–2 (1999).

2. C. Cenciarelli et al., Curr.Biol. 9, 1177–9 (1999).

3. D. Skowyra, K. L. Craig, M. Tyers, S. J. Elledge, J. W. Harper, Cell 91, 209–219 (1997).

4. D. E. Nelson, S. J. Randle, H. Laman, Open.Biol. 3, 130131 (2013).

5. S. K. Ganesh et al., Nat.Genet. 41, 1191–1198 (2009).

6. K. Ding et al., Mayo Clin.Proc. 87, 461–474 (2012).

7. H. P. van der et al., Nature 492, 369–375 (2012).

8. N. Soranzo et al., Nat.Genet. 41, 1182–1190 (2009).

9. S. J. Randle, D. E. Nelson, S. P. Patel, H. Laman, J.Pathol. 237, 263–272 (2015).

10. M. Lomonosov et al., PLoS.One. 6, e21165 (2011).

11. A. Di Fonzo et al., Neurology 72, 240–245 (2009).

12. F. R. Teixeira et al., Biochem.J. (2016).

13. H. Laman et al., Embo J 24, 3104–16 (2005).

14. H. J. Kuiken et al., J.Cell Mol.Med. 16, 2140–2149 (2012).

15. J. Kang and K. C. Chung, Cell Mol.Life Sci. 72, 181–195 (2015).

16. V. S. Burchell et al., Nat. Neurosci. 16, 1257–1265 (2013).

17. S. Vingill et al., EMBO J. (2016).

18. B. Fabre et al., Mol.Syst.Biol. 11, 771 (2015).

19. M. P. Bousquet-Dubouch et al., Mol.Cell Proteomics. 8, 1150–1164 (2009).

20. S. R. Stott et al., J. Pathol. (2019).

21. M. Bader et al., Cell 145, 371–382 (2011).

22. M. Bader, E. Arama, H. Steller, Development 137, 1679–1688 (2010).

23. R. C. Ballesteros et al., PLoS.One. 14, e0212481 (2019).

24. S. P. Patel, S. J. Randle, S. Gibbs, A. Cooke, H. Laman, Cell Mol.Life Sci. (2016).

25. e. K. Meziane, S. J. Randle, D. E. Nelson, M. Lomonosov, H. Laman, J.Cell Sci. 124, 2175–2186 (2011).

26. C. G. Mullighan et al., Nature 446, 758–764 (2007).

27. K. Okuyama et al., PLoS.Genet. 15, e1008280 (2019).

28. R. Somasundaram, M. A. Prasad, J. Ungerback, M. Sigvardsson, Blood 126, 144–152 (2015).

29. T. Inaba et al., Science 257, 531–534 (1992).

30. R. P. Kuiper et al., Leukemia 21, 1258–1266 (2007).

31. M. P. Kamps, C. Murre, X. H. Sun, D. Baltimore, Cell 60, 547–555 (1990).

32. E. Coyaud et al., Blood 115, 3089–3097 (2010).

33. M. Busslinger, N. Klix, P. Pfeffer, P. G. Graninger, Z. Kozmik, Proc.Natl.Acad.Sci.U.S.A 93, 6129–6134 (1996).

34. S. Iida et al., Blood 88, 4110–4117 (1996).

35. Z. Kozmik, S. Wang, P. Dorfler, B. Adams, M. Busslinger, Mol.Cell Biol. 12, 2662–2672 (1992).

36. Y. Miyazaki, X. Sun, H. Uchida, J. Zhang, S. Nimer, Oncogene 13, 1721–1729 (1996).

37. Y. Liu et al., Mol.Cell Biol 26, 3114–3123 (2006).

38. H. Kishi et al., Blood 99, 576–583 (2002).

39. K. Weston, Nucleic Acids Res. 20, 3043–3049 (1992).

40. G. A. Miranda et al., Mol.Immunol. 38, 1151–1159 (2002).

41. K. Larsen and C. Bendixen, Mol.Biol.Rep. 39, 1517–1526 (2012).

42. Nature 489, 57–74 (2012).

43. Y. Linderson et al., EMBO Rep. 5, 291–296 (2004).

44. D. Fitzsimmons et al., Genes Dev. 10, 2198–2211 (1996).

45. R. Hannah, A. Joshi, N. K. Wilson, S. Kinston, B. Gottgens, Exp.Hematol. 39, 531–541 (2011).

46. J. Malaterre et al., Stem Cells 26, 173–181 (2008).

47. P. Urbanek, Z. Q. Wang, I. Fetka, E. F. Wagner, M. Busslinger, Cell 79, 901–912 (1994).

48. J. E. Lee et al., Science 268, 836–844 (1995).

49. J. A. Nielsen, J. A. Berndt, L. D. Hudson, R. C. Armstrong, Mol.Cell Neurosci. 25, 111–123 (2004).

50. Y. Zhang and B. D. Manning, Cell Cycle 14, 2011–2017 (2015).

51. P. A. Li, X. Hou, S. Hao, J.Neurosci.Res. 95, 2025–2029 (2017).

52. J. Steffen, M. Seeger, A. Koch, E. Kruger, Mol.Cell 40, 147–158 (2010).

53. H. Digaleh, M. Kiaei, F. Khodagholi, Cell Mol.Life Sci. 70, 4681–4694 (2013).

54. W. T. Chang, H. I. Chen, R. J. Chiou, C. Y. Chen, A. M. Huang, Biochem.Biophys.Res.Commun. 334, 199–206 (2005).

55. J. J. Gasiorek and V. Blank, Cell Mol.Life Sci. 72, 2323–2335 (2015).

56. Y. Tsuchiya et al., Mol.Cell Biol. 31, 4500–4512 (2011).

57. S. Koizumi, J. Hamazaki, S. Murata, Proc.Jpn.Acad.Ser.B Phys.Biol.Sci. 94, 325–336 (2018).

